# AlveolEye: Rapid and precise lung morphometry guided by computer vision

**DOI:** 10.1101/2025.09.30.679300

**Authors:** Joseph Hirsh, Samuel Hirsh, Shawyon P. Shirazi, James Hirsh, Imani Douglas, Shriya Garg, Yeongseo Son, Arline Pierre-Louis, Claire Bunn, Christopher S. Jetter, Taiana J. James, Alexandria L. Sharkey, John T. Benjamin, Jonathan A. Kropski, Jennifer M. S. Sucre, Nicholas M. Negretti

## Abstract

Rigorous and reproducible evaluation of lung tissue under different conditions is necessary to interpret development, injury, and pharmacologic interventions. Common histological measurements in the distal lung include mean linear intercept (MLI) as a metric of alveolarization and airspace volume density (ASVD) as a metric of airspaces relative to tissue. Historically, these have been performed manually in a time-intensive process, with reproducible trends, but a high degree of variability between individuals. To improve the reproducibility and throughput of lung morphometry, we developed AlveolEye, an open source, semi-automated, computer vision-assisted tool that rapidly and reproducibly calculates MLI and ASVD from images of standard hematoxylin and eosin (H&E) stained tissue sections. AlveolEye-assisted MLI calculation closely aligns with manually-derived measurements for corresponding images, with preservation of trends in measurements between non-injured controls and neonatal mice subjected to two different injury models. Analyzing human tissue of varying ages suggests that the approach developed in AlveolEye is generalizable across species. Notably, AlveolEye markedly reduced the average variation across individual analyzers, with the greatest improvement in precision among individuals with the least experience in performing lung morphometry. The design of AlveolEye is intentionally semi-automated, preserving the investigator’s ability to assess and adjust parameters based on sample characteristics. AlveolEye facilitates efficient lung morphological measurements on larger sample sizes, allowing for greater statistical power for preclinical studies, and improves precision across individual observers, allowing for improved rigor in experimental design and execution.

## Introduction

In lung research, histological examinations allow for quantification and comparison of alveolar structure under different conditions, exposures, and models of lung injury^1^. Although qualitative differences among samples are observable between images, quantitative lung morphometry allows for measurement and comparison of multiple images and the usage of descriptive and inferential statistics. Quantification of imaging data is a critical part of the rigor with which these data are presented and interpreted, providing information about the consistency of the data obtained and assurance that presented images are representative of the aggregated data.

Despite automation of many laboratory techniques in data acquisition and analysis, lung morphometry remains a predominantly manual task. The most common metrics include a measurement of alveolar number and size (quantified by either radial alveolar count or mean linear intercept (MLI)) and the thickness of the walls between distal airspaces (quantified by airspace volume density, (ASVD)) ^2–4^. As commonly performed, lung morphometry is painstaking and time-consuming, with poor inter-rater precision ^5^. These logistical considerations undoubtedly constrain the sample size of model organisms used in experiments, with the unintended consequence of missing small effect sizes that require larger sample numbers for statistical power. Although the advent of machine learning has allowed for the development of fully automated systems of lung morphometry^5–8^, these approaches often require significant computer coding background to deploy and have not been widely adopted by the field.

To calculate lung morphometric measurements, such as MLI and AVSD, researchers examine the alveolar parenchyma and avoid airways and vessels in their analysis^1^. To do this, researchers rely on pattern recognition of airways and vessels for manual exclusion. Advances in computational technology have facilitated computer vision approaches whereby a large number of manually annotated images could be used to train neural networks to do the same pattern recognition of airways and vessels based on histologic features of these structures ^9^. Previously, generalizable pattern recognition algorithms applied to other imaging modalities such as computed tomography imaging have been used to perform segmentation of specific structures within other organs, providing the foundation for this study^10^.

To allow for high-throughput, reproducible lung morphometric data acquisition and analysis, we have developed a computer vision-assisted application that allows for human decision making at each stage of analysis, while improving consistency and reducing analysis time per image. Here we report the development of AlveolEye and compare AlveolEye with traditional manual image approaches by both trained experts (>1 year experience) and novices in lung morphometry (<1 year experience). AlveolEye is available as a plugin to the Napari image viewer and runs on individual personal computer systems, without the need for data upload or to rely on continued operation of web infrastructure. To facilitate longevity in the face of changing software ecosystems and extend future software capability, AlveolEye is open source and downloadable from www.sucrelab.org/alveoleye, with an overall goal to provide a resource to the lung research community that improves the rigor, precision, and efficiency of lung morphometry.

## Results

### A trained computer vision model identified airways and vessels in lung histology images to exclude them from morphometric analysis

A schematic of the AlveolEye image analysis pipeline can be found in **Figure 1**. In summary, we trained the computer vision model for 1000 iterations on 676 manually annotated hematoxylin and eosin (H&E) stained tissue images from control mouse lungs between P7 and P28. A MaskR-CNN^11^ computer vision model has an advantage of recognizing feature instances in an image and generating a mask (represented as a confidence map) that indicates which pixels compose each feature. AlveolEye can then calculate the mean linear intercept (MLI) and airspace volume density (ASVD) of the non-masked regions in each target image. AlveolEye calculates MLI by measuring the mean length of the line segments (chords) that lie along the test lines and span airspace. AlveolEye also returns ASVD defined as the percentage of all pixels representing the lung area that is airspace.

**Figure 1.**
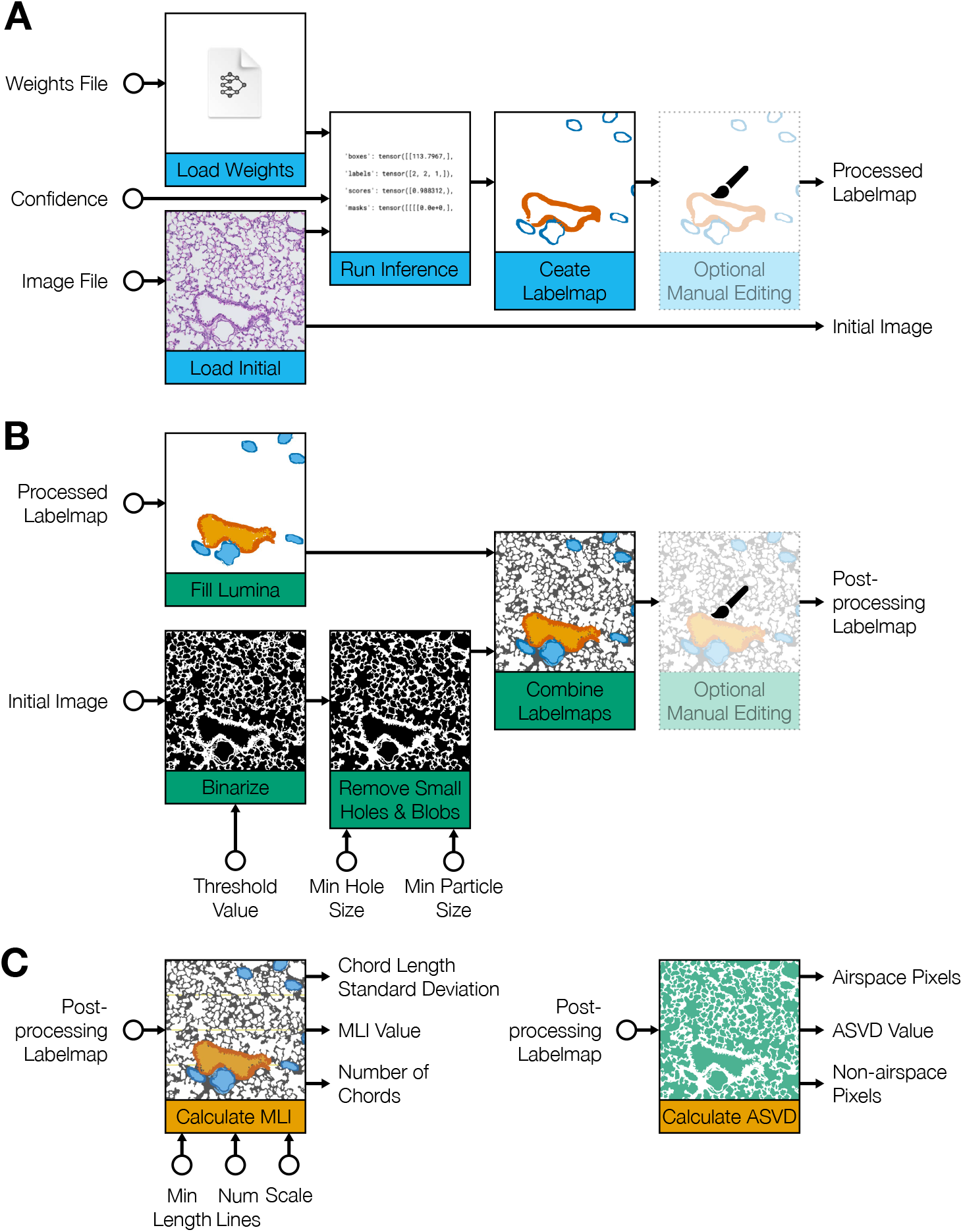
A deep learning neural network was combined with classical image analysis as a plugin to the Napari image viewer to facilitate review and analysis of all results. (**A**) The input image is processed with the trained computer vision model to identify and segment airway and vessel tissue; (**B**) AlveolEye employs classical image segmentation methods to identify the remaining image features, including airway and vessel lumens, airspace, and parenchyma. (**C**) AlveolEye performs morphometric assessments (MLI and ASVD) from the processed image.

The computer vision-assisted component of AlveolEye processes an image from an H&E-stained slide using the trained model, producing a map of the airway and vessel regions. To account for the visual differences between vessels and airways, two classifiers were trained. The image features used to predict airways were highly specific, only identifying airway tissue. The image features used for vessel prediction assigned high confidence values to both vessels and some airways, suggesting that the model extracted visual similarities between the two structures in some cases (**Figure 2**). AlveolEye combines both airway and vessel maps to flood-fill the airway and vessel lumens for exclusion, mitigating the impact of misattribution errors from either classifier. Tunable parameters such as computer vision confidence, background threshold, small particle removal, small gap removal, minimum line length, and image scale allow for the adjustment of AlveolEye to accommodate a variety of samples. Masking of cellular debris and other small particles and setting a minimum line length (**Figure 3**) to reduces the effect of tissue artifacts during MLI and ASVD calculation.

**Figure 2.**
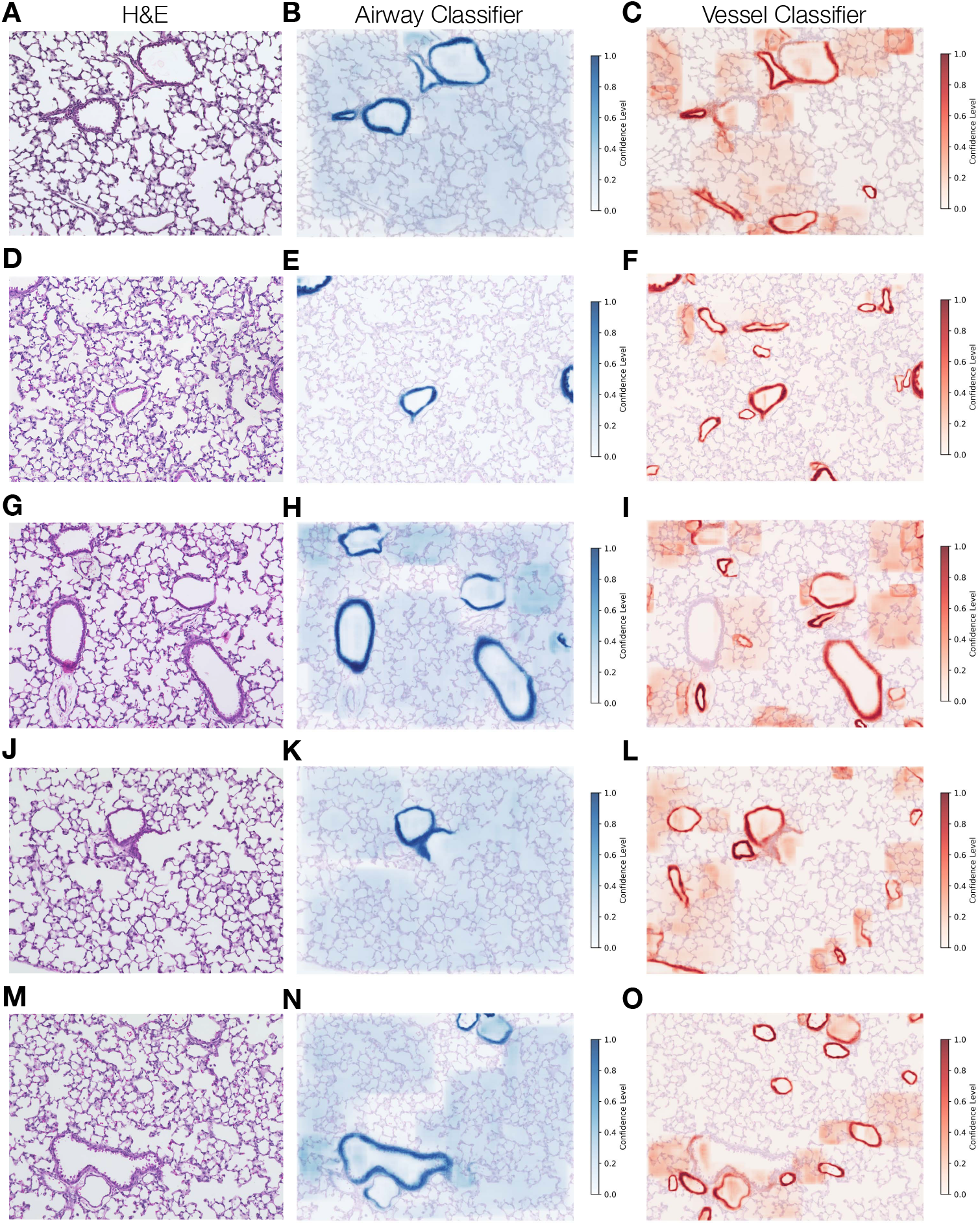
Two trained Mask R-CNN classifiers identify vessels and airways in lung histology images. To expedite the process of segmenting features to exclude from morphometric calculations, a computer vision-assisted model was trained on manually segmentized H&E-stained images (**A, D, G, J, M**), to identify airways (**B, E, H K, N**), and vessels (**C, F, I, L, O**). The output from the trained classifiers is displayed as a heatmap for airways (blue) and vessels (orange) overlayed on the original H&E-stained image. Darker colors indicate greater confidence of accurate classification by the computer vision-assisted model.

**Figure 3.**
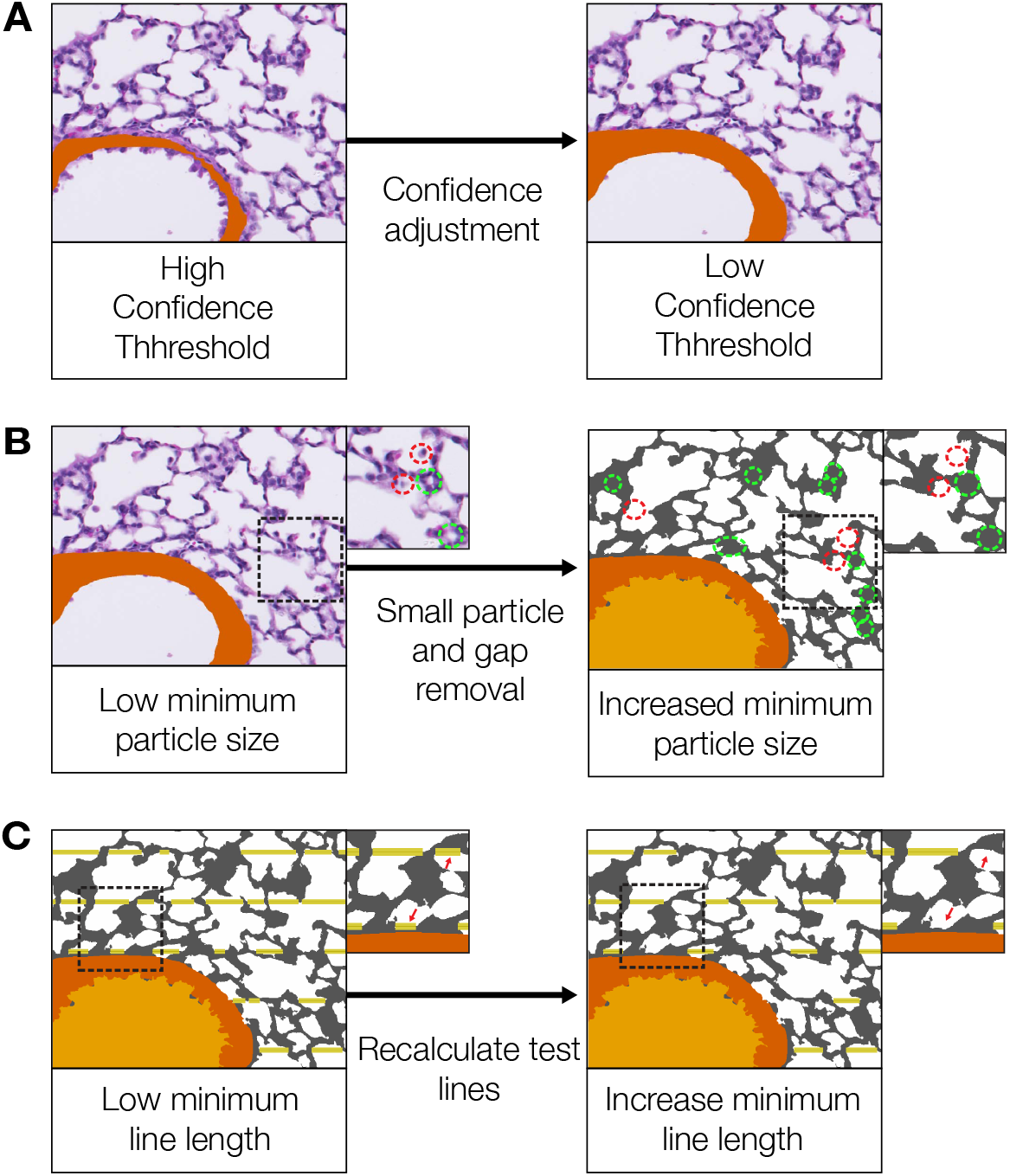
Parameters are adjustable to tune AlveolEye to different tissue conditions. The user adjustable parameters include: (**A**) Computer vision confidence threshold, where increasing the threshold makes the prediction more specific and less inclusive. Computer vision-based segmentation can also be disabled in favor of manual annotation. (**B**) Removal of small particles (red dotted outline), and subsequent of small gaps (green dotted outline) to account for artifacts of tissue thresholding and segmentation. (**C**) Minimum line length for the test lines used to measure mean linear intercept. Dotted lines indicate location of the magnified inset.

### AlveolEye reliably calculates lung morphometrics with values similar to human calculations

We assessed accuracy and reproducibility of AlveolEye morphometric quantification during development in postnatal day (P) 14 and P28 mice by comparing AlveolEye to manual image analysis performed by trained individuals of varying levels of experience (**Figure 4A, S2**). The MLI calculated MLI values calculated were generally consistent with manually measured MLI (**Figure 4B**). At P14, the average AlveolEye-assisted MLI value was 24.08 μm (SD = 2.21), compared to the average manually measured MLI value of 21.22 μm (SD = 1.33). By P28, the average AlveolEye-assisted MLI value was 22.64 μm (SD = 1.13), compared to the average manually measured MLI value of 24.69 μm (SD = 1.03). We also found that as an individual scientist with limited practice performed manual measurements over 189 images, the measurements became closer to the AlveolEye-assisted measurements (*p* = 0.000876, by linear regression) (**Figure 4C**).

**Figure 4.**
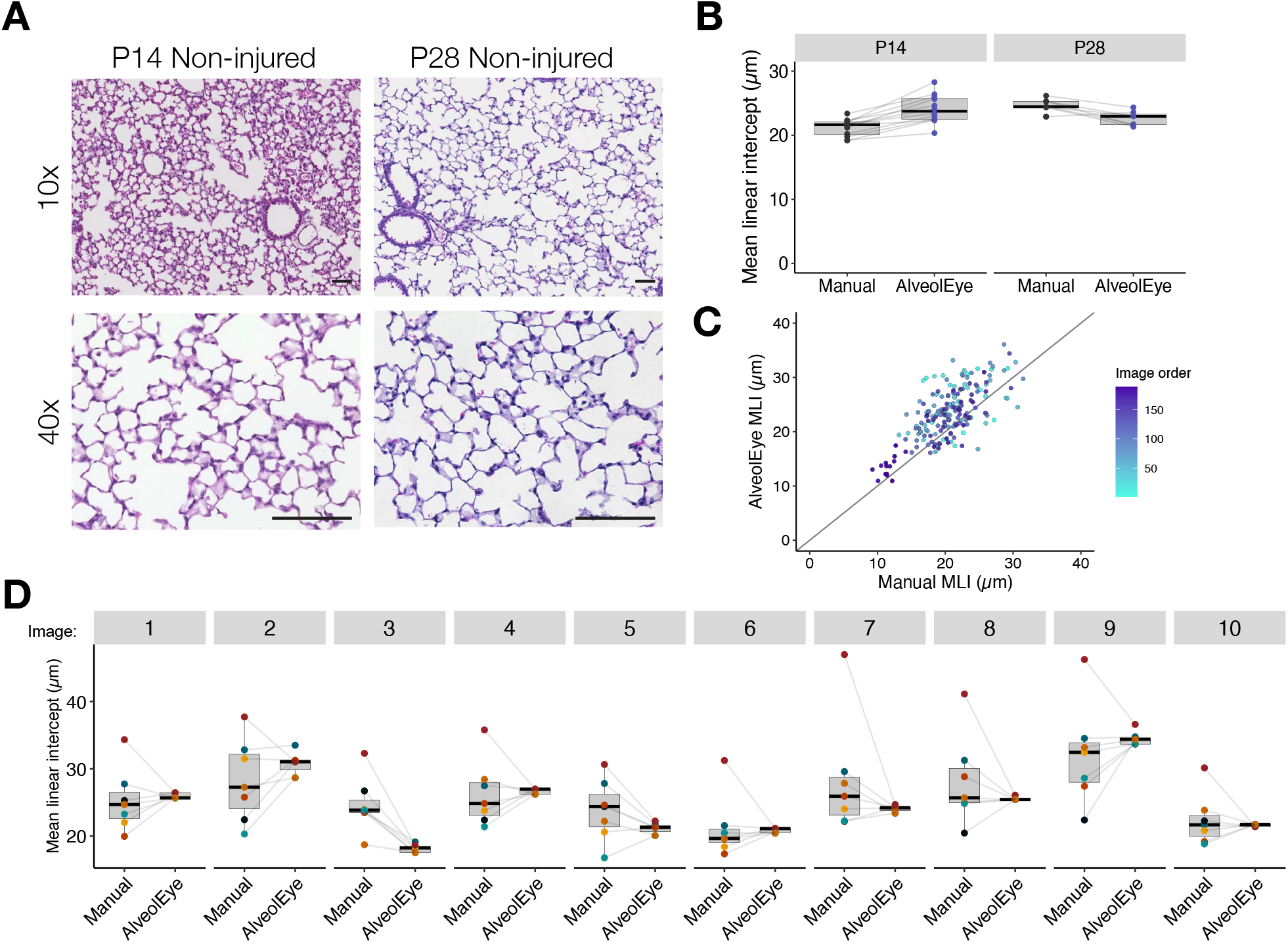
Calculation of MLI by AlveolEye for mice during alveologenesis closely aligns with manual MLI values with greater reproducibility between analyzers. (**A**) Representative images of H&E-stained lung sections of non-injured mice at P14 and P28 at 10x and 40x magnification. Scale bar = 100 μm. (**B**) Quantification of average MLI values calculated manually and by AlveolEye are similar for mice at P14 (*n* = 12) and P28 (*n* = 7). The box plots depict the median (center line) the 25^th^ and 75^th^ percentiles (box bounds), and whiskers which extend to 1.5 times the interquartile range. Each dot is the average MLI value from one animal. (**C**) Scatter plot with reference line (y = x) showing AlveolEye-assisted and manually derived MLI values colored by the order of image measurement, with later images in a darker color. Each dot represents the MLI of one image. (**D**) Comparison of multiple individuals measuring MLI manually and with AlveolEye across the same image set (n = 10 images), where each dot represents the measurement from one individual. Lines connect measurements made by the same individual (with analysis from additional images in Figure S2C).

We analyzed the variability of measurements by six individuals performing manual MLI calculations on 20 images from two non-injured mouse H&E-stained samples at P14. The manually derived MLI measurements obtained by 7 individuals has an average range of 14.16 μm (**Figure 4D**), with the highest variability coming from individuals with the least experience (defined as < 1 year) (**Figure S2A-C**). As the same individuals analyzed the same images with AlveolEye, the average range decreased to 1.93 μm (**Figure 4D, Figure S2A**). Two individuals tracked the time it took for manual analysis and AlveolEye-assisted analysis. AlveolEye reduced processing time by approximately 90%, from an average of 4 minutes 35 seconds per image for manual measurement, to an average of 25 seconds per image with AlveolEye assistance.

### AlveolEye is generalizable across different injury perturbations in mice

To assess whether AlveolEye could be used in the setting of pathology, we reanalyzed a previously published hyperoxia-exposed lung injury model.^4,12,13^ Mice were exposed to 85% oxygen from P2 through P14, and lung structure was compared to mice in normoxia (21%) at the same age. Consistent with previous published descriptions of this model, representative H&E-stained sections showed that hyperoxia exposure resulted in disrupted alveolarization, characterized by enlarged and simplified airspaces (**Figure 5A-B**). At P14, the average manually measured MLI for non-injured lungs was 22.26 μm (SD = 6.73) and 44.57 μm (SD = 8.54) for hyperoxia-exposed lungs. AlveolEye-assisted measurements demonstrated a similar trend, with average MLI values for 19.96 µm (SD = 4.87) for non-injured lungs and 37.29 µm (SD = 5.42) for hyperoxia-exposed lungs. At P21, we observed a similar trend, with the manually derived average MLI of 25.85 µm (SD = 1.89) for non-injured lungs and 47.56 µm (SD = 11.56) for hyperoxia-exposed lungs. AlveolEye-assisted values were 21.13 µm (SD = 1.78) in non-injured lungs and 53.10 µm (SD = 10.25) in hyperoxia-exposed lungs. We further assessed the AlveolEye-assisted MLI measurements with a second previously published lung injury model^14^. In this model, mice were exposed to 70% oxygen from P1 through P5 with intranasal LPS administration on P3 and P4. Representative H&E-stained sections demonstrated that this lung injury model also led to enlarged and simplified alveolar structures (**Figure 5C-D**). In this dataset, the average manually measured MLI in non-injured mice was 23.92 µm (SD = 1.40) compared to the average AlveolEye assisted measurement of 22.55 µm (SD = 1.05). For the injured mice, the average manual MLI was 30.83 µm (SD = 3.00) compared to the AlveolEye assisted measurement of 30.15 µm (SD = 2.63).

**Figure 5.**
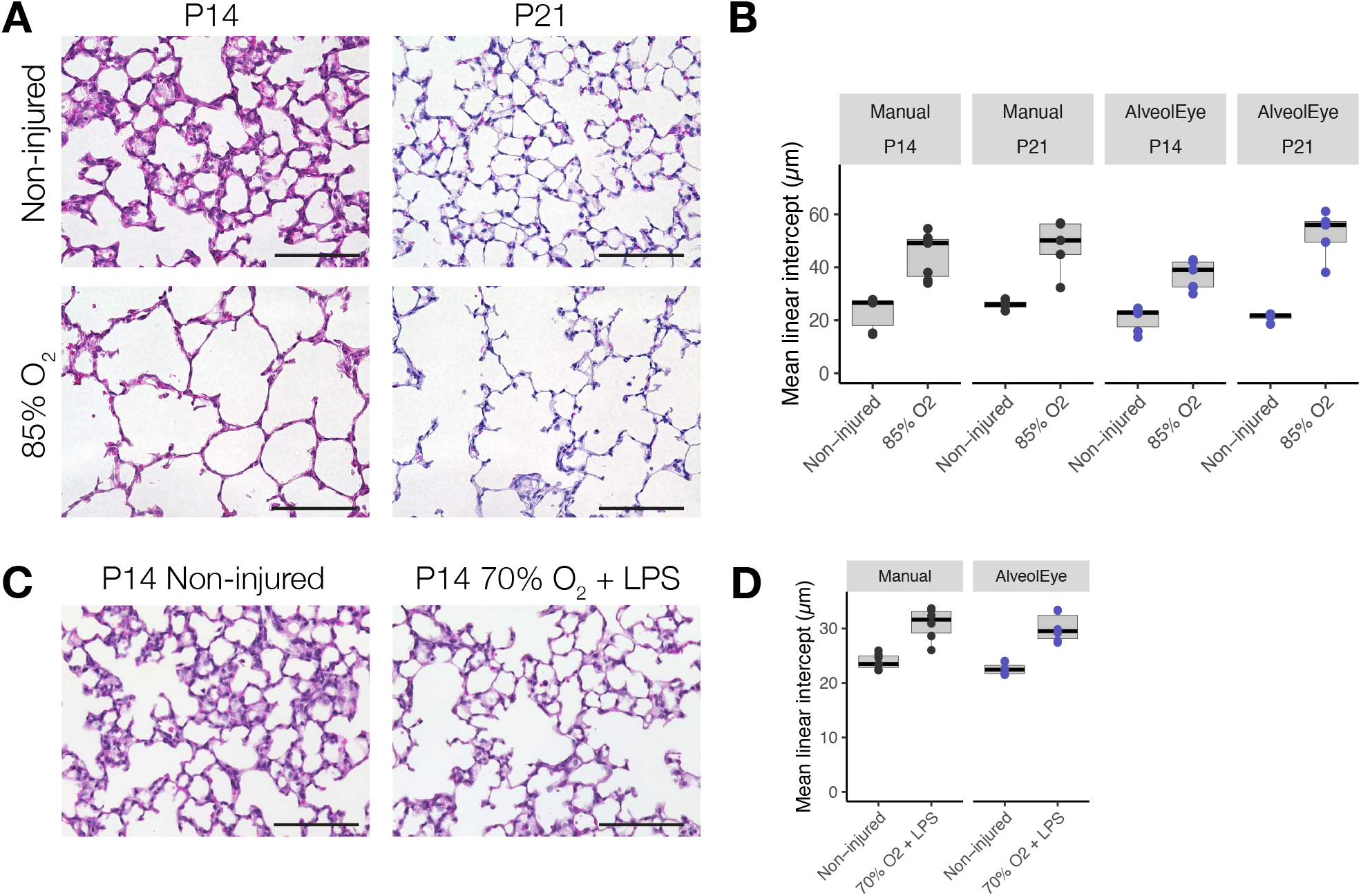
AlveolEye effectively quantifies structural changes after hyperoxia lung injury models in mice. (**A**) Representative H&E-stained lung sections from non-injured control and hyperoxia-induced mice at P14 and P21. Scale bar = 100 µm. (**B**) Manually measured and AlveolEye calculated MLI measurement in non-injured and hyperoxia-exposed mice at P14 and P21, with box plots depicting the median (center line), 25^th^ and 75^th^ percentiles (box bounds), and whiskers which extend to 1.5 times the interquartile range, and where each dot represents the average of the images for one mouse. (**C**) Representative H&E-stained lung sections from non-injured and 70% O2 +LPS for P1-5 injured mice and analyzed at P14. (**D**) Manual and AlveolEye quantification of MLI, with box plots depicting the median (center line), 25^th^ and 75^th^ percentiles (box bounds), and whiskers which extend to 1.5 times the interquartile range. Each dot represents the average from10 images per mouse.

### AlveolEye effectively assists measuring MLI on human lung images

We applied AlveolEye to lung tissue from eight human autopsy samples without any reported primary lung disease. The 10 H&E-stained images per sample (n = 7 samples, 70 images total) showed human lung alveolar structures as larger in size and with lower alveolar density compared to mice, consistent with prior reports^15^ (**Figure 6-B, Figure 4A-B**). Despite the inherent variability between mice and human samples, AlveolEye yielded comparable results to manual measurements, despite having not been trained on human tissue. To improve adaptability across species and injury models, we designed AlveolEye with tunable parameters (e.g. computer vision confidence, background threshold, small particle removal, small gap removal, minimum line length and image scale). To accommodate the difference in human and mouse lung morphology, we adjusted these parameters at the level of individual samples (preserving the same settings across images from the same slide). As expected, using unadjusted default parameters yielded more measurements less consistent with manually-obtained MLI, (**Supplementary Figure 3**). This highlights the usefulness of semi-automated analysis tools, that allow for tuning based on sample features. In addition to analyzing AlveolEye on adult human samples, we tested AlveolEye on five neonatal lung samples. In the AVSD measurements performed on 42 total images, manual analysis was comparable to AlveolEye (**Figure 6C-D**).

**Figure 6.**
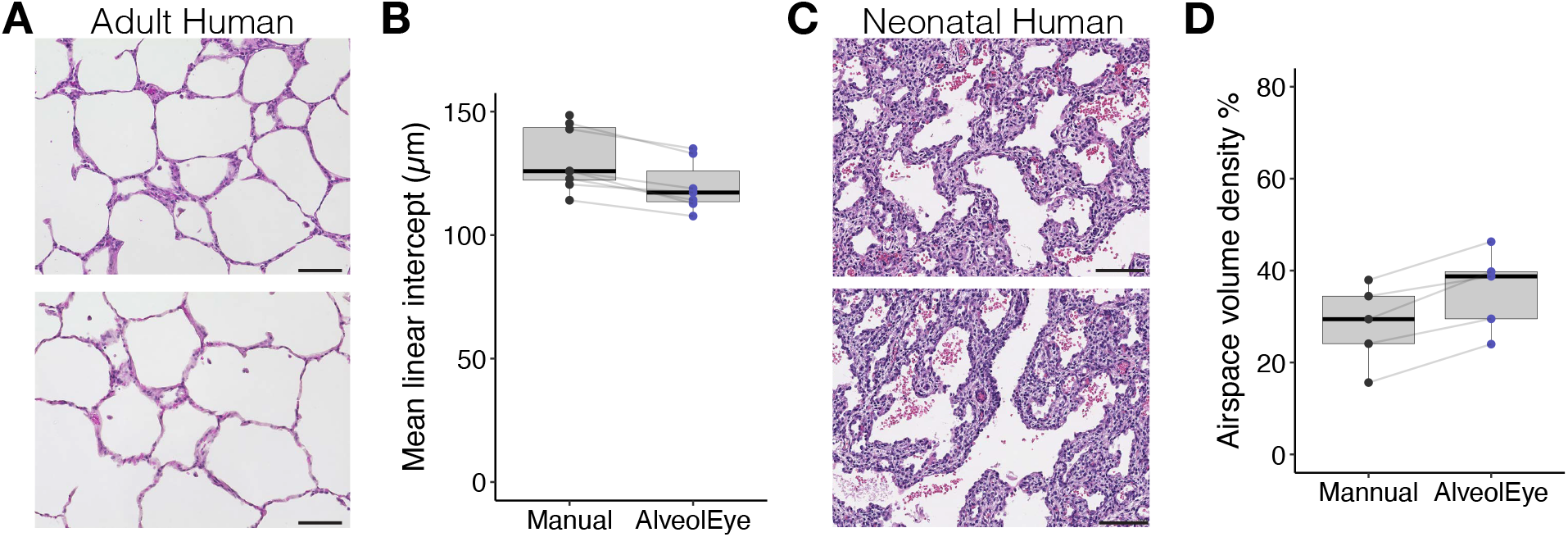
Despite being trained on mouse lungs, AlveolEye performs effectively on human lung tissue. (A) Visual representation of representative H&E-stained adult human lung images used for manually derived and AlveolEye-assisted MLI measurements. Scale bar = 100 µm. (B) Comparison of the average manually-measured MLI values to AlveolEye across 7 human lung samples, where each dot represents the average from 10 images from individual samples. (C) Representative H&E-stained human neonatal lung samples used for ASVD measurements. Scale bar = 100 µm. (D) Comparison of the average manual and AlveolEye assisted ASVD measurements across 5 neonatal lung samples, where each dot represents the average from 7-10 images from individual samples. Box plots depict the median (center line), 25^th^ and 75^th^ percentiles (box bounds), and whiskers which extend to 1.5 times the interquartile range.

### The impact of the number of lines on the mean linear intercept measurement was assessed with AlveolEye

There is no gold standard for the number of test lines in MLI. To assess the relationship between variance and line length, we compared MLI calculations with 3 randomly placed lines on each with 10 randomly placed lines on the same set of 11 images. The three-line MLI calculations resulted in an average variance of 8.26 µm, (range 2.95-18.08 µm) (**Figure 7A**). The 10-line MLI calculations resulted in an average variance of 2.76 μm, (range 0.79 - 5.54 μm) (**Figure 7B**). With increasing the number of test lines (up to 50), we found that variance decreased as the number of lines increased across all images and stabilizes (< 2 μm from the mean) starting at around ten lines (**Figure 7C**).

**Figure 7.**
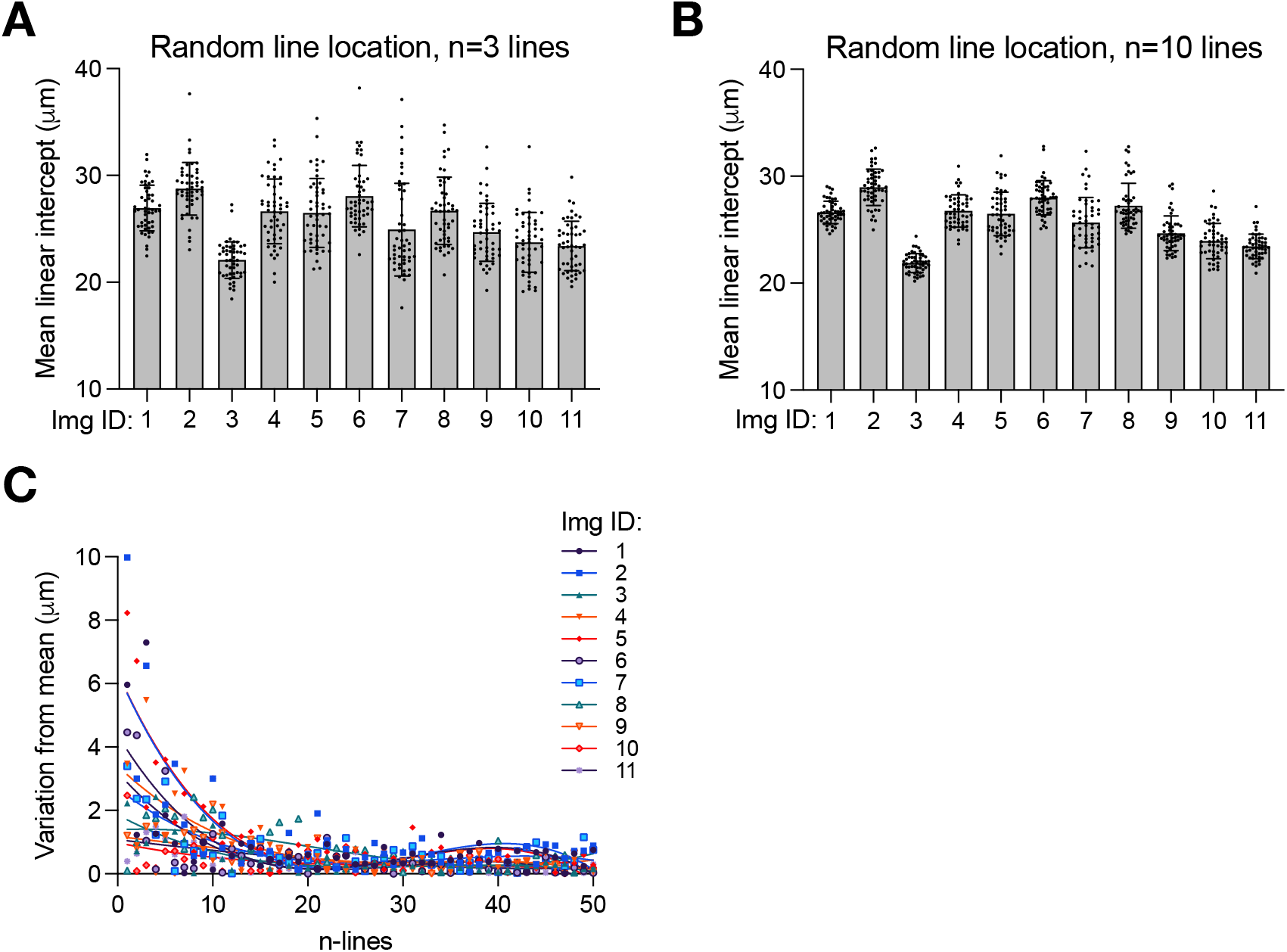
Increasing the number of test lines decreases the impact of random variation of line placement. Eleven images were evaluated. (A) MLI values on 11 images with 3 test lines and random line locations. Dots represent the MLI value from each of the 50 trials. (B) MLI values on the same 11 images with 10 test lines and random line locations. Dots represent the MLI value from each of the 50 trials. (C) The variation of the MLI from the mean (μm) as a function of the number of test lines on each of the images (color) where each dot represents the MLI from each image at the given number of lines.

## Discussion

AlveolEye-assisted measurements enhance reproducibility, precision, and speed of lung morphometric analysis. We found that AlveolEye reproduces manually-calculated trends of relatively increased MLI measurements in non-injured vs injured lungs and does so with far less inter-rater variance when compared to manually methods. Our analysis uncovered new insights into the MLI metric itself, such as the impact of the number and location of test lines. Taken together these data support the use of AlveolEye to add rigor, reproducibility, and efficiency to studies of lung development, injury, and repair.

AlveolEye is not the only effort to automate lung morphometric analysis, with prior work incorporating aspects of machine learning to a greater or lesser degree^10,16,5^. Examples include a machine-learning tool using a U-Net neural network to provide an important foundation for subsequent morphometric automation^5^, a MATLAB-based automated morphometric calculation^17^, and most recently, Deep-Masker, an AI tool that measures chord length in murine models of COPD^18^. To these foundational studies, AlveolEye adds significant advances, including being constructed as open-source software to allow for adaptability to future platforms and technologies and. In addition, AlveolEye was designed with lung biologists as the end-user, requiring little to no background in coding or computer science on the part of the user to launch and operate. Our goal was to create a flexible application for analyzing lung tissue in variable stages of lung development, injury and repair. Indeed, an arguable advantage of AlveolEye over entirely automated platforms is that it requires the user to scrutinize every image for parameter tuning, thus ensuring a layer of human oversight for quality control and consistency.

Overall, AlveolEye reduces variance when compared to manual calculation and decreases the time required for morphometric analysis, with the biggest improvement in consistency observed in individuals with least experience with morphometric analysis. Because of variability within and between individuals performing manual calculation, all MLI for a set of comparison experiments are often done by a single individual, requiring repetition of the entire dataset with addition of new samples or conditions. Our data also suggest that humans become more proficient with practice, meaning that the variability from even a single individual is not applied equally across a dataset, with images analyzed earlier having a greater variability than those analyzed later. By contrast, AlveolEye remains consistent throughout use. In manual analysis, the location (and number) of the guide test lines is another source of variability across published methods and research groups. The increase from 3 to 10 lines with manual MLI would more than triple the time required per image, but with AlveolEye, the computational time required is essentially the same for all the images regardless of line number, allowing for a more accurate representation of approximated alveolar count, with no loss in efficiency.

Because AlveolEye is consistent, its biases are also consistent. However, when used to compare control and perturbed conditions (e.g., injury, gene manipulations, treatment), systematic biases of AlveolEye unrelated to tissue structure will affect all experimental groups evenly. The user can also mitigate systemic errors by adjusting parameters and intervening when needed (**Figure 3A-C**). Notably, AlveolEye does not perform well on samples with incomplete H&E staining and/or excessive cellular accumulation in the alveolar spaces in the cases of hemorrhage or alveolar exudative processes. To overcome some of these limitations, we have implemented a feature that allows for removal of small particles/cells, which is effective for cells in airspaces. While this approach still fails for large clumps of debris, the user can also remove unwanted regions manually. Furthermore, AlveolEye does not account for adventitia around large vessels, which may need manual annotation if large vessels are captured in the image field. Future training and augmentations such as blur and color jittering would likely improve the model’s ability to handle cases of incomplete staining or hemorrhage, although for cases of severe pulmonary hemorrhage, lung morphometry is perhaps not the appropriate analytical tool. The hemorrhage itself prevents lungs from being physiologically inflated at the time of fixation, a requirement for a morphometric approach.

Presently, AlveolEye only performs MLI and ASVD, two among many possible lung morphometry calculations ^1^. The open-source nature of AlveolEye and its modular software architecture make it possible for future extension to include additional metrics, e.g., radial alveolar counts and alveolar septal tip length, among others. With rapid advances in computational sophistication, we envision that AI-based tools will grow more suited to aid researchers in tedious tasks. As is the case with AlveolEye, we hope that the augmentation of sample number capacity and rigor does not supplant the opportunity for novel pattern identification and discovery. With AlveolEye decreasing the practical barriers for image analysis, there is an opportunity for larger experimental replicates and expanded sampling of different spatial areas of the lung, generating more nuanced but uniformly robust quantification.

## Methods

### Animal sample collection and care

All mice were on the C57BL/6 background under a 12:12 hr dark/light cycle with free access to food and water. Mice were either exposed to 85% oxygen from P2 until P14 as described previously^19^, or were exposed to 70% oxygen from P1 until P5 with intranasal LPS instillation on P3 and P4 (1 ug/g) as described previously^14^. Control mice were housed in room air conditions. Lung tissue was collected by inflation and fixed by gravity filling with 10% buffered formalin at P14, P21, or P28 and paraffin embedded.

### Human subjects and samples

The Human Infant Lung Repository at Vanderbilt University Medical Center has been reviewed and approved by the Vanderbilt University Institutional Review Board with a non-human subjects determination (no additional consent required). All samples are de-identified with the exception of sex, gestational age at birth, age at time of death, cause of death.

### Manual calculation of mean linear intercept (MLI)

H&E-stained images were captured on a Keyence BZX-800 with either a 20x or 40x objective. Manual MLI measurements were calculated with ImageJ, on images with the appropriate scale set. For each image, three evenly spaced horizontal lines were drawn across the lung tissue as guides to measure intercepts. Lines were drawn across alveolar airspaces avoiding large airways, blood vessels, and artifacts. The length of the lines was averaged for each image. For measurements representing a mouse, at least 10 images were measured and then averaged.

### Manual airspace volume density (ASVD) calculation

Manual ASVD measurements were calculated in ImageJ after setting the appropriate scale. For each image, airspaces were selected using the ‘magic wand tool’ with a tolerance of 10. The total combined area of the selected airspaces was divided by the total pixel area to create a ratio.

### Training Dataset Creation

The training dataset comprised 676 images of hematoxylin and eosin-stained (H&E) lung slides. Training annotation masks for each of the 676 H&E-stained images were created using Adobe Photoshop (versions 22.0.0 to 24.0.1). Each mask is an annotated version of a histological image in which all parenchyma pixels are blue, vessel pixels are green, airway pixels are yellow, and non-tissue spaces, blood cells, artifacts, and other non-tissue elements are white. To ensure consistent and efficient annotation, we leveraged Photoshop Actions to create a uniform, semi-automated workflow. We document the workflow steps in detail in the Supplementary Methods.

### AlveolEye Segmentation, processing, and calculations

AlveolEye used a Mask R-CNN (v0.16) model from the torchvision^20^. Python package with a ResNet-50 FPN^21^ backbone and pre-trained COCO_V1 weights. The torchvision FastRCNNPredictor and MaskRCNNPredictor were tuned to accommodate three output classes: airway epithelium, vessel endothelium, and background. We trained the computer vision model on a Nvidia RTX4000 GPU. The program augments a random 15% of the images on-the-fly during training. The possible augmentations were horizontal and vertical flips, color jittering, random rotations, Gaussian blur, and affine transformations. We trained the classifiers for 1,000 epochs with each image in the training set appearing twice per epoch.

The Mask R-CNN model predicts the presence of two classes, airway epithelium and vessel endothelium, in provided histological images. A feature prediction consists of probability scores for each pixel, representing the likelihood that the pixel belongs to the respective class. For each class, the program compares the pixel probabilities to a user specified threshold to determine which pixels to assign the relevant class label. The program overlays the binarized class-specific label maps to create a combined 2D semantic label map. Each pixel is classified as background, airway epithelium, or vessel endothelium.

AlveolEye then uses classical image processing to segment the parenchyma and alveoli, remove small gaps in the tissue and blobs in the airspace, and identify lumina within the vessels and airways to exclude from further calculations. All image processing operations use OpenCV’s image analysis capabilities. After converting the input image to grayscale, a threshold value is determined using Otsu’s method to create a binary image mask. AlveolEye adds 20 to the threshold value, because we observed that this augmentation improves segmentation quality. Small particles are removed by labeling connected components smaller than a user-defined size as background. To remove small gaps, the same operation is performed on the inverse image. AlveolEye then segments airway and vessel lumina based on the labeled tissue regions in their respective label maps by calculating the contour around the labeled mask and filling the enclosed region.

To calculate mean linear intercept, AlveolEye overlays a specified number of evenly spaced horizontal test lines atop the image, excluding segments of the test lines that overlap with non-airspace pixels. AlveolEye then computes the mean chord length for all line segments. To calculate airspace volume density, AlveolEye counts the number of pixels in the processed image that correspond with alveolar airspaces. This value is divided by the total number of pixels in the processed image that corresponds to parenchyma plus airspaces, resulting in a fraction of lung tissue which is airspace.

More detailed information on each computational step can be found in the supplementary materials.

### Statistics and plots

Boxplots and scatterplots were generated in R (version 4.5.0) using ggplot2. Linear regressions were done in R. Bar plots were generated in Graphpad Prism. Values in the plots either represent the average from an individual animal, or individual image, as indicated in the figure legends.

## Code and data availability

All relevant code related to this manuscript and the AlveolEye plugin are available on GitHub: https://github.com/SucreLab/AlveolEye/.

## Acknowledgements

This work was supported by the Francis Family Foundation (JAK, JMSS, NMN), R01HL168556 (JMSS), K08HL143051 (JMSS), and U01HL175444 (JAK, JMSS). This material is based upon work supported by the National Science Foundation Graduate Research Fellowship Program under Grant No. DGE-2146755 (YS). Any opinions, findings, and conclusions or recommendations expressed in this material are those of the author(s) and do not necessarily reflect the views of the National Science Foundation. We thank Alexander Dean for assistance capturing images for the training set. We are grateful to Christopher Wright, *DPhil* and Vivian Siegel, PhD for their thoughtful discussions about this project.

## References

1. Hsia, C.C.W., Hyde, D.M., Ochs, M., and Weibel, E.R. (2010). An Official Research Policy Statement of the American Thoracic Society/European Respiratory Society: Standards for Quantitative Assessment of Lung Structure. Am J Respir Crit Care Med 181, 394–418. 10.1164/rccm.200809-1522ST.

2. Pérez-Bravo, D., Myti, D., Mižíková, I., Pfeffer, T., Surate Solaligue, D.E., Nardiello, C., Vadász, I., Herold, S., Seeger, W., Ahlbrecht, K., et al. (2021). A comparison of airway pressures for inflation fixation of developing mouse lungs for stereological analyses. Histochem Cell Biol 155, 203–214. 10.1007/s00418-020-01951-0.

3. Litzlbauer, H.D., Neuhaeuser, C., Moell, A., Greschus, S., Breithecker, A., Franke, F.E., Kummer, W., and Rau, W.S. (2006). Three-dimensional imaging and morphometric analysis of alveolar tissue from microfocal X-ray-computed tomography. American Journal of Physiology-Lung Cellular and Molecular Physiology 291, L535–L545. 10.1152/ajplung.00088.2005.

4. Sucre, J.M.S., Vickers, K.C., Benjamin, J.T., Plosa, E.J., Jetter, C.S., Cutrone, A., Ransom, M., Anderson, Z., Sheng, Q., Fensterheim, B.A., et al. (2020). Hyperoxia Injury in the Developing Lung Is Mediated by Mesenchymal Expression of Wnt5A. Am J Respir Crit Care Med 201, 1249–1262. 10.1164/rccm.201908-1513OC.

5. Salsabili, S., Lithopoulos, M., Sreeraman, S., Vadivel, A., Thébaud, B., Chan, A.D.C., and Ukwatta, E. (2021). Fully automated estimation of the mean linear intercept in histopathology images of mouse lung tissue. J Med Imaging (Bellingham) 8, 027501. 10.1117/1.JMI.8.2.027501.

6. Pu, J., Leme, A.S., de Lima E Silva, C., Beeche, C., Nyunoya, T., Königshoff, M., and Chandra, D. (2023). Deep-Masker: A Deep Learning-based Tool to Assess Chord Length from Murine Lung Images. Am J Respir Cell Mol Biol 69, 126–134. 10.1165/rcmb.2023-0051MA.

7. Madi, A., Politis, D.A., Salsabili, S., and Chan, A.D.C. (2025). Automated mean linear intercept measurement: quantifying bias and parameter sensitivity in lung morphometry. Physiol Meas 46. 10.1088/1361-6579/adf0bd.

8. Politis, D., Salsabili, S., and Chan, A.D.C. (2022). An Automated Tool to Assess Air Space Size in Histopathology Images of Lung Tissue. In 2022 IEEE International Instrumentation and Measurement Technology Conference (I2MTC), pp. 1–6. 10.1109/I2MTC48687.2022.9806556.

9. Rayed, Md.E., Islam, S.M.S., Niha, S.I., Jim, J.R., Kabir, M.M., and Mridha, M.F. (2024). Deep learning for medical image segmentation: State-of-the-art advancements and challenges. Informatics in Medicine Unlocked 47, 101504. 10.1016/j.imu.2024.101504.

10. Garcia-Uceda, A., Selvan, R., Saghir, Z., Tiddens, H.A.W.M., and de Bruijne, M. (2021). Automatic airway segmentation from computed tomography using robust and efficient 3-D convolutional neural networks. Sci Rep 11, 16001. 10.1038/s41598-021-95364-1.

11. He, K., Gkioxari, G., Dollár, P., and Girshick, R. (2018). Mask R-CNN. Preprint at arXiv, https://doi.org/10.48550/arXiv.1703.06870 10.48550/arXiv.1703.06870.

12. Ambalavanan, N., Nicola, T., Hagood, J., Bulger, A., Serra, R., Murphy-Ullrich, J., Oparil, S., and Chen, Y.-F. (2008). Transforming growth factor-beta signaling mediates hypoxia-induced pulmonary arterial remodeling and inhibition of alveolar development in newborn mouse lung. Am J Physiol Lung Cell Mol Physiol 295, L86–95. 10.1152/ajplung.00534.2007.

13. Ramani, M., Bradley, W.E., Dell’Italia, L.J., and Ambalavanan, N. (2015). Early Exposure to Hyperoxia or Hypoxia Adversely Impacts Cardiopulmonary Development. Am J Respir Cell Mol Biol 52, 594–602. 10.1165/rcmb.2013-0491OC.

14. Shirazi, S.P., Negretti, N.M., Jetter, C.S., Sharkey, A.L., Garg, S., Kapp, M.E., Wilkins, D., Fortier, G., Mallapragada, S., Banovich, N.E., et al. (2025). Bronchopulmonary dysplasia with pulmonary hypertension associates with semaphorin signaling loss and functionally decreased FOXF1 expression. Nat Commun 16, 5004. 10.1038/s41467-025-60371-7.

15. Irvin, C.G., and Bates, J.H. (2003). Measuring the lung function in the mouse: the challenge of size. Respiratory Research 4, 1. 10.1186/rr199.

16. Khanna, A., Londhe, N.D., and Gupta, S. (2023). A Deep Attention-based U-Net for Airways Segmentation in Computed Tomography Images. Curr Med Imaging 19, 361–372. 10.2174/1573405618666220630151409.

17. Sallon, C., Soulet, D., Provost, P.R., and Tremblay, Y. (2015). Automated High-Performance Analysis of Lung Morphometry. Am J Respir Cell Mol Biol 53, 149–158. 10.1165/rcmb.2014-0469MA.

18. Pu, J., Leme, A.S., de Lima E Silva, C., Beeche, C., Nyunoya, T., Königshoff, M., and Chandra, D. (2023). Deep-Masker: A Deep Learning-based Tool to Assess Chord Length from Murine Lung Images. Am J Respir Cell Mol Biol 69, 126–134. 10.1165/rcmb.2023-0051MA.

19. Sucre, J.M.S., Vickers, K.C., Benjamin, J.T., Plosa, E.J., Jetter, C.S., Cutrone, A., Ransom, M., Anderson, Z., Sheng, Q., Fensterheim, B.A., et al. (2020). Hyperoxia Injury in the Developing Lung Is Mediated by Mesenchymal Expression of Wnt5A. Am J Respir Crit Care Med 201, 1249–1262. 10.1164/rccm.201908-1513OC.

20. TorchVision maintainers and contributors (2016). TorchVision: PyTorch’s Computer Vision library. (GitHub).

21. Ren, S., He, K., Girshick, R., and Sun, J. (2016). Faster R-CNN: Towards Real-Time Object Detection with Region Proposal Networks. Preprint at arXiv, https://doi.org/10.48550/arXiv.1506.01497 10.48550/arXiv.1506.01497.

