## Supplementary Information for "AlveolEye: Rapid and precise lung morphometry guided by computer vision"

### *Supplementary Figures*

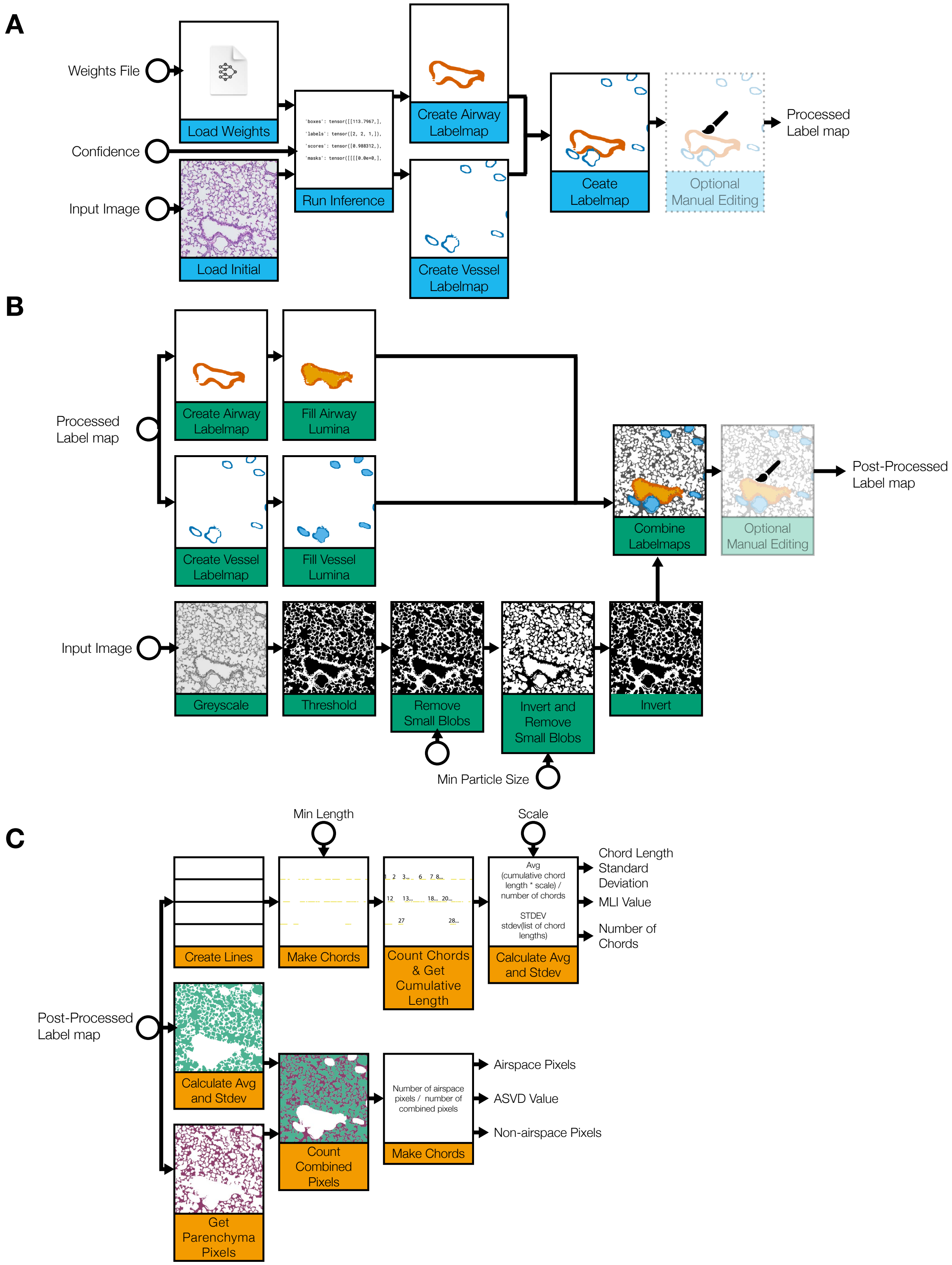

**Figure S1. Detailed diagram of image processing steps.** (A) The input image is processed with the trained computer vision model to identify and segment airway and vessel tissue; (B) AlveolEye employs classical image segmentation methods to identify the remaining image features, including airway and vessel lumens, airspace, and parenchyma. (C) AlveolEye performs morphometric assessments (MLI and ASVD) from the processed image. Circles indicate user input points.

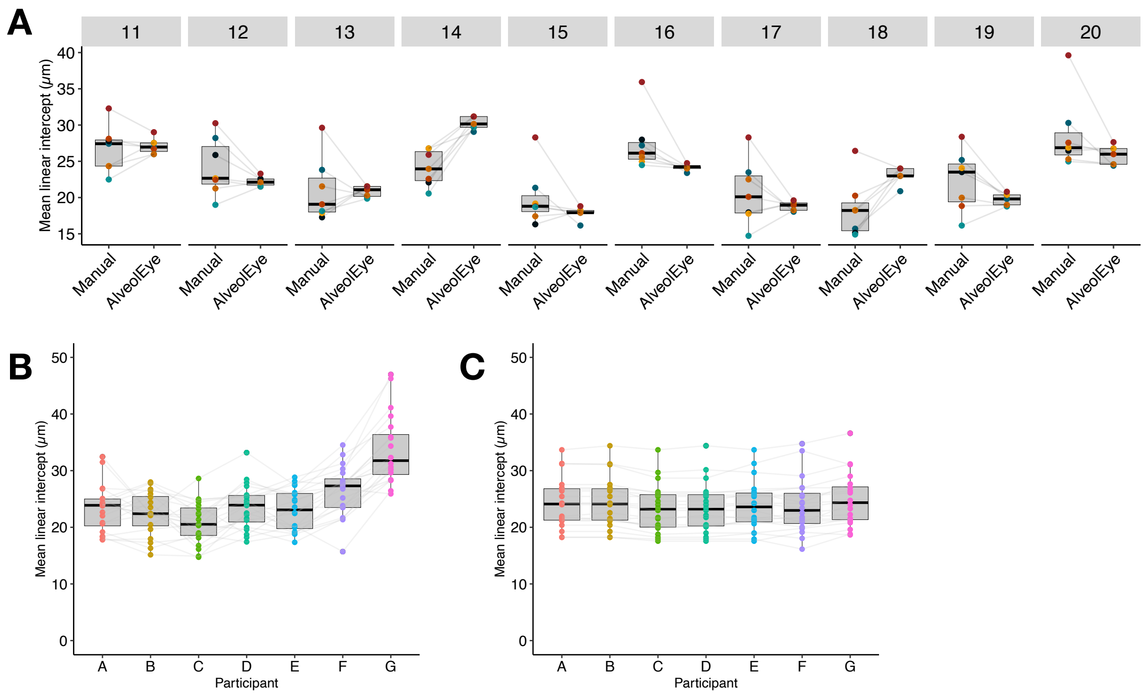


**Supplementary Figure 2**. AlveolEye reduces variability among individuals when compared to manual measurements. (A) Comparison of multiple individuals measuring MLI manually and with AlveolEye across the same image set (n = 10 additional images), where each dot represents the measurement from one individual. Lines connect the same individual. (B) Comparison of multiple individuals measuring MLI manually and with (C) AlveolEye across the same image set, where each dot represents an image. In panels B and C, participants are sorted by experience, from most to least experience. Lines connect the same image, n = 20 images.

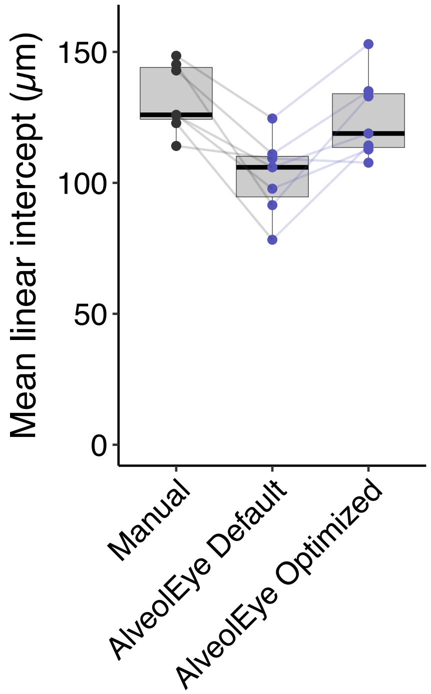


**Supplementary Figure 3**. Application of default parameters to diverse samples results in inaccurate measurements. However, optimizing parameters improves measurement consistency with manual images.

### Supplementary Methods

#### *Mask Creation Workflow*

1. Step 1: An automated action (1) thresholds the histological image (with 174 as the threshold value) to color all dark pixels (tissue, blood cells, artifacts) black and the background white and (2) selects all black pixels and colors them blue. We automated this step to ensure consistency, efficiency, and accuracy across histological images, and to reduce human effort and error. We chose the value 174 because it produced the best thresholding for most images. We determined this threshold value through visual inspection and subjective evaluation. Frequently during annotation, we adjusted the action threshold level to better suit a specific image when the default value captured either too many or too few pixels.
2. Step 2: The user uses the "Magic Wand" tool to select all blue pixels corresponding to tissue, effectively excluding blood cells, artifacts, and other non-tissue elements. A future improved workflow could replace manual selection with an action to exclude blood cells by incorporating a size-based filter that ignores pixel blobs smaller than a set minimum acceptable size. AlveolEye employs this method in the post-processing stage.
3. Step 3: An automated action (1) inverses the selection to select all non-tissue pixels, (2) colors all non-tissue pixels white, and (3) inverses the selection to re-select all tissue pixels. This process is well suited for automation because it consists of a precise sequence of operations that require no human judgment.
4. Step 4: The user uses the pencil tool to annotate all vessel tissue green and airway tissue yellow. Because annotations differ entirely from one image to another, we could not automate this process in Photoshop; AI processing in AlveolEye *is* the automated version of this step.

#### ***Computer Vision Training***

AlveolEye uses Mask R-CNN model within torchvision[9] with a ResNet-50 FPN[10] backbone and pre-trained COCO_V1 weights. The FastRCNNPredictor and MaskRCNNPredictor from torchvision are tuned to accommodate the output classes: airway epithelium, vessel endothelium, and background.

We trained AlveolEye on a Nvidia RTX4000 GPU. Training images are loaded using PIL and converted to PyTorch[11] tensors. We then construct two DataLoaders (batch_size=10, num_workers=0) that shuffle the training image set but not the validation set. The program augments a random 15% of the images using the torchvision.transforms.v2 module on-the-fly during training. The possible augmentations are horizontal and vertical flips, color jittering, random rotations, Gaussian blur, and affine transformations.

We trained the classifiers for 1,000 epochs using SGD (learning rate=0.005, momentum=0.9, weight_decay=0.0005) on gradients calculated by the backpropagation algorithm. The model trained on each image twice per epoch (n_repeat_images=2). We used a StepLR scheduler to decay the learning rate by 0.1 every 20 epochs. Each epoch, we computed training loss and per-class classification/box-regression metrics, ran the validation loader to obtain mean average precision (mAP) and related detection metrics, and logged all scalar values plus the first batch of raw images (training and validation) to TensorBoard. We saved intermediate model checkpoints every 50 epochs and the final model at epoch 1,000. Once training completes, the program saves the model weights into a .pth file.

#### ***Segmentation Processing***

AlveolEye performs image segmentation using a torchvision implementation of a Mask R-CNN model (v0.16) with ResNet-50 FPN backbone. AlveolEye initializes the model immediately after the user loads a trained weights file through the graphical user interface (GUI). If the user does not provide a trained weights file, AlveolEye loads the model with the default trained weights when the user initializes processing with the “Process” button. The first time the user selects the default trained weights, AlveolEye downloads the trained weights file from Google Drive using gdown. After the model loads, AlveolEye replaces the default classification and mask prediction heads with custom layers configured for three classes: airway epithelium, vessel endothelium, and parenchyma.

AlveolEye reads the input image with the Python Imaging Library (PIL), converts the image to RGB (if not in RGB format already), and transforms the image via torchvision into a normalized float32 tensor. If a CUDA-compatible GPU is available, the program moves the model and image tensor from the CPU to the GPU. The Mask R-CNN then performs inferencing with the loaded model. The output is a list of instance-level predictions that include binary masks, class labels, bounding boxes, and confidence scores.

A label map-generation function ingests the model output and produces the final semantic label map for the processing step. To build the final label map, the program generates separate airway and vessel label maps through identical processes: It iterates through every model prediction and saves instances in which (1) the class matches the class being processed and (2) the confidence score exceeds the confidence threshold provided by the user through the GUI. This process produces a list of NumPy arrays, one for each instance of the target class. AlveolEye stacks the 2D arrays to create a 3D array where each layer corresponds to a separate mask and the depth axis indexes each instance. A combined binary mask for all instances of the class is then generated by checking each pixel position across all stacked instance masks to see if any predict that the pixel belongs to the class. If at least one mask contains a nonzero value at that location, the corresponding pixel in the combined mask is marked as True; otherwise, it is False.

AlveolEye then creates an empty label map with the same shape as the input image. It assigns the airway epithelium label to each pixel in the final label map for which the corresponding pixel in the combined airway binary mask is True; it then assigns the vessel endothelium label to each pixel in the final label map for which the corresponding pixel in the combined vessel binary mask is True (potentially overwriting airway labels where the airway and vessel masks overlap). This process produces a 2D semantic label map in which each pixel is classified as background, airway epithelium, or vessel endothelium.

***Image Post-processing***

The post-processing step ingests the initial image and the model-generated instance-level predictions and produces a single, cleaned, semantically segmented label map; distinct labels represent airway epithelium, vessel endothelium, airway lumen, vessel lumen, parenchyma, and alveoli. All image processing operations use OpenCV's image analysis capabilities.

AlveolEye first converts the initial histologic image to grayscale. It then thresholds the grayscale image using either a user-defined threshold value (if manual thresholding is enabled through the interface) or Otsu's method. Thresholding binarizes the image by assigning a pixel value of zero to the "background" and one to the "foreground." AlveolEye adds 20 to the threshold value computed by Otsu's method; we observed that this correction improves segmentation quality. The program removes small particles by labeling connected components smaller than a user-defined size as background. To remove small holes, it inverts the image to treat holes as connected components and applies the same size filtering operation. The program then inverts the image back to its original foreground-background orientation.

The thresholded label map and the final processing label map are then used to create the complete post-processing label map. The program first generates intermediate label maps to isolate specific classes: It creates a parenchyma label map by assigning the parenchyma label to all pixels at positions where the corresponding thresholded label map pixel is one and assigning the alveoli label to all pixels at positions where the corresponding thresholded label map pixel is zero. Then, AlveolEye generates the airway epithelium and vessel endothelium label maps by assigning the respective class label to all pixels in which the same pixel in the processing label map is that class; it assigns zero to every other pixel.

AlveolEye then segments airway and vessel lumina based on the labeled tissue regions in their respective label maps. AlveolEye uses the airway epithelium and vessel endothelium label maps to generate a binary mask for each in which pixels assigned the respective class label are set to one, and all other pixels are set to zero. AlveolEye computes the contours of the binary mask. Contours are the minimum-size continuous outer boundaries surrounding each connected region of epithelial tissue. The program also calculates a bounding box for each contour and checks whether any side lies within a specified distance from the image edge. If the bounding box is sufficiently close to an edge, the tissue is assumed to be truncated by the border and unclosed. To close the open region, the program draws a line along the intersecting image border. The program then recalculates the contours to reflect the updated boundaries. For each contour, the program generates a filled mask of the enclosed region and computes its centroid. The program then locates the connected component in the thresholded image that contains the centroid and assigns the appropriate lumen label to all pixels within that component.

#### *Automated Morphometry Metric Calculations*

AlveolEye can calculate MLI and airspace volume density (ASVD) by analyzing the fully-segmented post-processed label map. The MLI calculation demands user inputted parameters: number of lines, minimum length, and scale (hereafter bracketed for readability). The ASVD calculation does not require user inputted parameters.

#### *Mean Linear Intercept*

The program overlays [number of lines] evenly-spaced horizontal test lines atop the post-processed label map. The program excludes segments of the test lines that overlap with non-airspace pixels. These non-airspace pixels are all pixels not labeled with the alveoli label (seven by default) in the post-processed label map. The program excludes other segments that are shorter than [minimum length]. The remaining segments overlap with airspace and exceed the [minimum length]. We denote these included segments "chords." The "Assessments" layer depicts all chords.

The program computes the mean chord length: It calculates the combined length of all chords (A pixels). It counts the total number of chords (B chords). To calculate the unscaled MLI, it divides the cumulative chord length by the total number of chords (A pixels / B chords = C pixels per chord). To convert this average to alternative units, such as microns, the program multiplies it by [scale] (C pixels per chord * [scale] units per pixel = D units per chord). The resulting value (D units per chord) is the MLI.

In addition to computing the MLI, the program calculates the standard deviation of the chord lengths. The graphical widget displays the MLI, standard deviation, and total number of chords.

***Airspace Volume Density***

The program counts the number of pixels (A pixels) in the post-processed label map that correspond with alveoli. These alveoli pixels are all pixels labeled with the alveoli label (seven by default) in the post-processed label map. The program also counts the number of pixels (B pixels) in the post-processed label map that correspond with parenchyma. These parenchyma pixels are all pixels labeled with the parenchyma label (six by default) in the post-processed label map. The program sums the number of alveoli pixels with the number of parenchyma pixels to acquire the total number of alveoli and parenchyma pixels (A pixels + B pixels = C pixels). The total number of alveoli and parenchyma pixels equals the number of pixels in the image that do not correspond with airway epithelium, vessel endothelium, airway lumen, vessel lumen, or the blocking label.

The program divides the number of alveoli pixels by the total number of alveoli and parenchyma pixels (A pixels / C pixels = D). To acquire a percentage representation, the program multiplies the resulting fraction by 100 (D * 100 = E). The result (E) is the ASVD. This process, in contrast to the MLI calculation, does not correspond with any layer in the viewer.
